# Coding region changes in the Hippo pathway component YAP1 contribute to body size differences among mammalian species

**DOI:** 10.64898/2026.10.29.762931

**Authors:** Twinkle Bansal, Qingwei Zhu, Kunxin Luo, Iswar K. Hariharan

## Abstract

Animal species differ dramatically in their overall size, yet the mechanisms underlying these differences are poorly understood. The Hippo pathway, first discovered in *Drosophila*, and widely conserved in mammals, regulates the nuclear levels of YAP1, a transcriptional co-activator that enhances the expression of growth-promoting genes. While experimental manipulations of this pathway elicit dramatic effects on tissue growth and organ size, it is not known whether, and to what extent, evolutionary divergence of this pathway has contributed to size differences across animals. Here we show that although YAP1 protein sequence is highly conserved between diverse mammals such as humans, elephants and blue whales, it diverges considerably among rodent species. While the human YAP1 protein is very similar to that of the largest rodent, the capybara *Hydrochoerus hydrochaeris*, those of smaller rodents such as the mouse *Mus musculus* and the vole *Microtus oregoni* show considerable divergence. An increased ratio of non-synonymous versus synonymous substitutions (dN/dS) in clades of small rodents compared to the background is indicative of relaxed constraint in these lineages, suggesting relative positive selection. When expressed in the developing *Drosophila* wing, capybara YAP1 was more potent at promoting growth than that of the mouse or vole. Following transfection into mammalian cells, capybara YAP1 is more effective at promoting expression of a reporter gene than either mouse or vole YAP1. Thus, by combining computational and experimental approaches, we show that evolutionarily divergent coding-region changes in YAP1 have likely contributed to the evolution of smaller body size in clades of rodents.

## INTRODUCTION

J. B. S. Haldane began his famous essay “On being the right size” (Haldane, 1926), with the sentence “The most obvious differences between different animals are differences of size, but for some reason the zoologists have paid singularly little attention to them”. In the century since that essay was written, there has been much descriptive work on how some animals evolved to differ in size, but in most cases we still have a poor understanding of the functional genetic differences that are responsible for differences in size among animals. The gastrulating embryos of different species are remarkably similar in their size, yet the final sizes of organisms that develop from those embryos can differ dramatically (O’Farrell, 2004). From first principles, these size differences could arise from either differences in the rate of growth after gastrulation, the duration of growth, or a combination of both processes (Alberch et al., 1979). At least for those organisms that do not exhibit indeterminate growth (Hariharan et al., 2015), there are likely mechanisms, yet to be discovered, that sense the size of a growing organism and arrest growth at an appropriate final size for that species.

The pathway composed of growth hormone (GH), its receptor (GHR), and their downstream effectors, the insulin-like growth factors (IGF1 and IGF2) have been implicated in size differences across various species. Increased production of growth hormone can result in increased human size e.g. in acromegaly (Ranke and Wit, 2018). Conversely, mutations in the growth hormone receptor have been identified in isolated human populations that have short stature (Guevara-Aguirre et al., 2011). Mutations in the IGF1 locus have also been shown to be responsible for size differences in dogs and other canids (Sutter et al., 2007; Plassais et al., 2022). Additionally, several genes in the GH/IGF axis appear to have undergone positive selection in clades of giant cetaceans (Silva et al., 2023). Most recently, the Neanderthal GHR was found to have multiple alterations compared to most modern humans that make it more active in tissue culture assays, and that this variant persists in some modern humans via introgression (Kanis et al., 2026).

De novo, nonsynonymous mutations that alter the stability or function of the encoded proteins could result in body size changes. Alternatively, they could also occur in regulatory regions of genes with an effect on gene expression. Since the protein coding sequences of chimpanzees and humans are nearly identical, yet there are many obvious morphological differences, (King and Wilson, 1975) hypothesized that most of these morphological differences were the result of changes in regulatory sequences that impacted gene expression. Subsequently, the hypothesis that morphological changes arise primarily from mutations in cis-regulatory sequences gained ascendancy (Carroll, 2008). However, this notion has been challenged by others who question the idea that mutations affecting morphology should primarily involve non-coding sequences (Hoekstra and Coyne, 2007). Indeed, differences in body size between dogs is the result of changes to both non-coding and coding sequences. A large-effect mutation between large and small breeds is a single-nucleotide polymorphism (SNP) in a non-coding antisense RNA, IGF1-AS, which overlaps the IGF1 primary transcript and alters blood IGF1 levels (Plassais et al., 2022). Additionally, in some miniature breeds, there is a non-synonymous SNP in the IGF1R gene that results in an arginine to histidine change in the cysteine-rich ligand-binding domain in the receptor (Hoopes et al., 2012). Thus both coding region and regulatory changes in this pathway have occurred during dog evolution to affect body size.

Genetic screens in model organisms such as *Drosophila* have been extremely successful in identifying genetic pathways that alter growth during development. Among these is the Hippo pathway (Fu et al., 2022; Phillips et al., 2024), which is highly conserved among diverse taxa from insects to mammals (Sun et al., 2025). At the core of the pathway, are two kinases - Hippo (MST1/2 in mammals) and Warts (LATS1/2 in mammals) - that function in series to phosphorylate the transcriptional co-activator Yorkie/YAP, leading to its exclusion from the nucleus. Higher levels of nuclear Yorkie/YAP result in increased expressions of pro-growth and anti-apoptotic genes (**Figure 1A**). Experimentally altering the activity of this pathway can elicit dramatic alterations in tissue growth resulting in alterations in organ size (Dong et al., 2007). This led to the attractive notion that the Hippo pathway could function like a rheostat to regulate organ size. However, this idea has been questioned by experiments that abolish the entire output of the pathway (both positive and negative) that cause little effect during normal development, leading to an alternative hypothesis that the pathway functions only during ectopic growth (Kowalczyk et al., 2022). Findings that alterations in the Hippo pathway during evolution have a direct contribution to size differences between species would support a role for this pathway in determining the size of organs and organisms during development. Yet, to our knowledge, no systematic investigations of Hippo pathway genes in this context have been published. One study described a nonsynonymous SNP in YAP1 in large cetaceans (Sun et al., 2025), but since its predicted effect would be to reduce growth, it could not have contributed to the increased size of those lineages. Indeed, the authors hypothesized that this substitution might have evolved because it reduced cancer risk in these large long-lived mammals.

**Figure 1.**
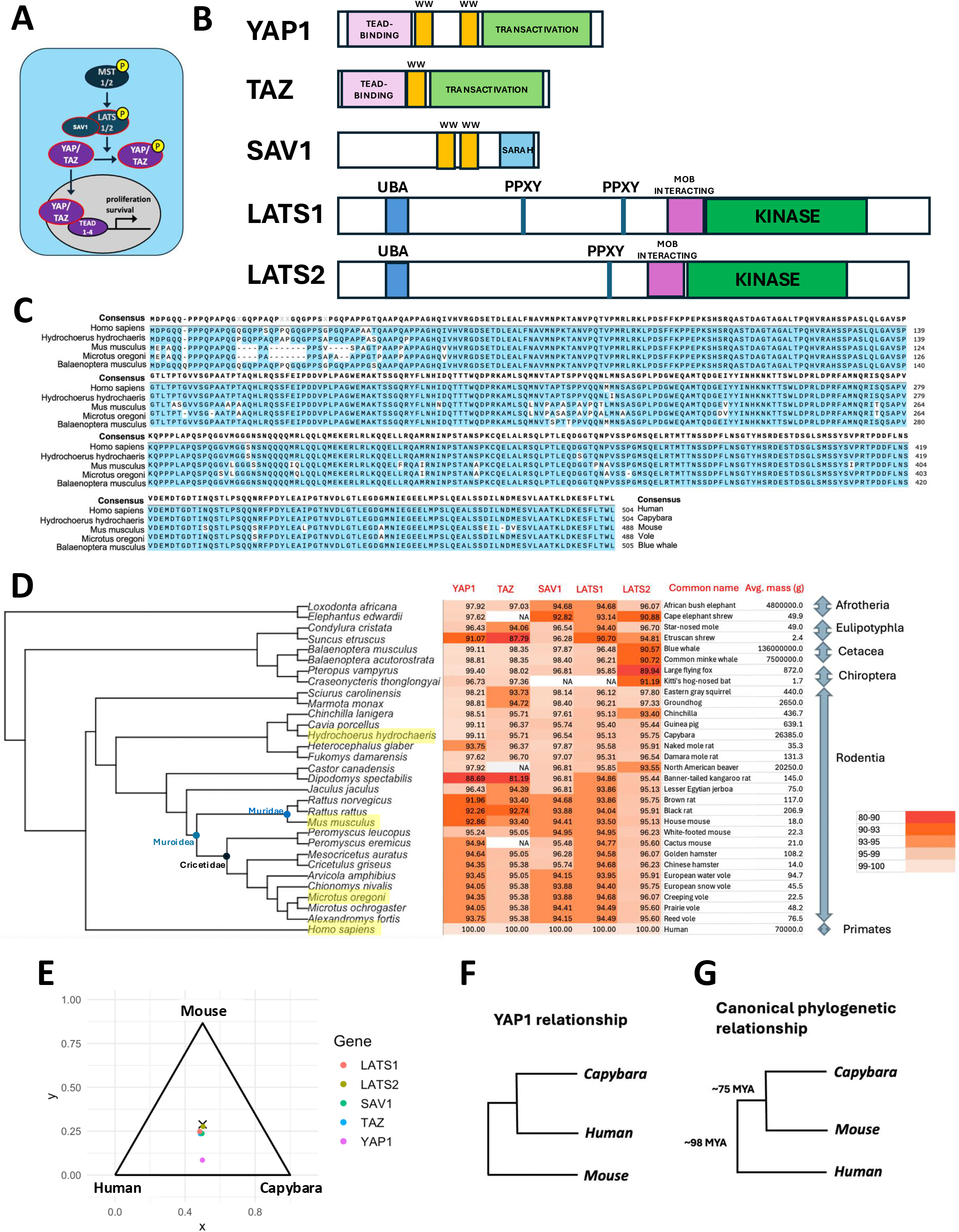
Divergence of YAP1 sequences in rodent clades. **A)** The mammalian Hippo pathway showing the upstream kinases of the core pathway, MST1 and MST2, the downstream kinases LATS1 and LATS2, and the scaffolding protein SAV1. Phosphorylation of the transcriptional coactivators YAP1 and TAZ by LASTS1/2 excludes them from the nucleus. In the nucleus YAP1 or TAZ bind to one of TEAD1-4 and promote the transcription of growth-promoting and anti-apoptotic genes. **B)** Schematic representations of the five proteins examined in this study, YAP1, TAZ (also known as WWTR1), SAV1, LATS1, and LATS2. Locations of key domains are indicated. The proteins and their domains are drawn to scale. **C)** Alignment of amino acid sequences of the YAP1 protein from *Homo sapiens* (Human), *Hydrochoerus hydrochaeris* (Capybara), *Mus musculus* (House mouse), *Microtus oregoni* (Creeping vole), and *Baelenoptera musculus* (Blue whale). Amino acids that match the consensus are shaded in blue, while those that are different are shown with a white background. **D)** A phylogeny showing 22 rodent species and humans. Outgroups included are Chiroptera (bats), Cetacea (whales and dolphins), Eulipotyphla (hedgehogs, shrews and moles), and Afrotheria (elephants and the elephant shrews). Common names and average mass are included. Shading indicates percentage identity among the trimmed sequences. **E)** Barycentric plot of human, mouse and capybara YAP1, TAZ, SAV1, LATS1, and LATS2. Displacement towards one side indicates increased sequence similarity between those two species. YAP1 human to capybara = 99.1%, Human to mouse = 92.8%, and mouse to capybara = 92.5%. **F)** Phylogenetic relationship based solely on YAP1 sequence **G)** Established phylogenetic relationship

Here we addressed this gap by examining the coding region of five genes encoding Hippo pathway components, SAV1, LATS1, LATS2, YAP1 and WWTR1/TAZ and found that while the YAP1 gene is highly conserved between mammals as diverse as humans and elephants, many nonsynonymous substitutions have occurred during the evolution of clades of small rodents. There is evidence of relaxed constraint, and potentially positive selection, occurring in these lineages. When assessed for their ability to promote growth in *Drosophila*, or activate a transcriptional reporter in mammalian cells, we found that the YAP1 genes of mice and voles were less effective than those of the human and the capybara.

## RESULTS

### YAP1 is highly conserved among mammals but shows divergence in rodent clades

We began our investigations by identifying mammalian lineages with coding region changes in five components of the Hippo pathway - the kinases LATS1 and LATS2, the adaptor protein SAV1 and the transcriptional co-activators YAP1 and WWTR1/TAZ (**Figure 1B**). We did this by performing pairwise alignments of sequences after matching isoforms from a range of placental mammals post-trimming of poorly aligned sequences (see Methods). This approach allowed us to quantify identities and substitutions in those regions that could be aligned with confidence. Using such an approach, we found that YAP1 is highly conserved between humans and phylogenetically distant mammalian species such as the African bush elephant *Loxodonta africana* and the blue whale *Balaenoptera musculus* (97.9% and 99.1% identity respectively) (**Figure 1C, D**). In contrast, we found that the YAP1 proteins encoded by several rodent species differed considerably, both from each other and from human YAP1. While the YAP1 protein encoded by the capybara *Hydrochoerus hydrochaeris*, a large rodent found in South America, shows a high degree of identity to the human protein (99.1%), the proteins from two smaller rodent species, the mouse *Mus musculus* and the vole *Microtus oregoni* are more divergent from the human sequence at the amino acid level (92.86% and 94.35% identity respectively). These alignments are shown in **Figure 1C**. This raised the possibility that YAP1 had undergone more changes in multiple species of small rodents.

Prompted by these findings, we used this approach to determine the pairwise sequence identity for five Hippo pathway genes between 31 different species including 22 rodent species. Here we show the identity of each of those species to their human counterparts (**Figure 1D**). Included in our analysis was a collection of 22 rodent species (order Rodentia), 2 species of Chiroptera (bats), 2 species of whales (Cetacea) and 2 representatives of the order Eulipotyphla, the star-nosed mole and the Etruscan shrew, which is the smallest known extant mammal by mass (Galán-Puchades et al., 2021). We took care to choose one relatively larger and one smaller species from each order/superorder outside of Rodentia. The group that is phylogenetically most distant to humans in our analysis is the Afrotheria, which is a superorder including the African bush elephant and its close and yet small phylogenetic relative, the Cape elephant shrew. From this analysis, it is evident that YAP1 in several clades of rodents is more different from the human protein than that from more distantly related groups such as Afrotheria and Cetacea. This is most obvious in the species from the Muroidea supergroup of rodents which includes rats, mice, hamsters and voles. Within this group, the most divergent YAP1 genes are from the Muridae family that includes mice and rats (92.0 - 92.9% identity to humans). Also within the Muroidea group are the Cricetidae which includes deer mice, hamsters and voles (93.8 - 95.2%). For rodents outside of the Muroidea family the YAP1 gene of the Banner-tailed kangaroo rat (88.7%) and the Naked mole rat (93.8%) show more divergence from human YAP1. Remarkably, in contrast to the YAP1 genes from small rodents that show many differences from the human, YAP1 from two relatively large rodents, the YAP1 proteins of the capybara (99.1%) and the North American beaver (97.9%) are much more similar to the human protein. Notably, outside of the rodents, the YAP1 protein has also diverged extensively from the human in the smallest of all mammals, the Etruscan shrew. That YAP1 genes from diverse large mammals are extremely similar while those of smaller mammals have diverged raises the possibility that some of these changes might contribute to reducing the extent of growth in these small mammals, possibly by reducing YAP1 activity.

Similar comparisons were conducted with the other four genes from the Hippo pathway, the YAP1 paralog TAZ, the adapter protein SAV1 and the kinases LATS1 and LATS2 (**Figure 1D**). TAZ genes from rodents do not exhibit the same divergence from other mammalian sequences that YAP1 does. Rather, there are a few species from different taxa where TAZ has diverged more. These include the Black rat, the Banner-tailed kangaroo rat and the Etruscan shrew. For SAV1 and LATS1, many species of rodents have more divergent genes. However, these differences are less than for YAP1 and there are not obvious differences in divergence between large rodents and small rodents. For LATS2, more divergence is apparent in more distant orders including Chiroptera (bats) and Cetacea (whales) as well as the Cape elephant shrew though not the other species from the Afrotheria group, the African bush elephant. Notably, for the Etruscan shrew, which is the smallest extant mammal by mass, four different Hippo pathway genes (YAP1, TAZ, LATS1 and LATS2) show more differences than for most other pairwise comparisons. Thus a divergence among rodent lineages and a difference between small and large rodents is most obvious with respect to YAP1.

To illustrate the relative similarity of the genes from three species, the human, the capybara, and the mouse, we used a barycentric plot where the three species are at the apices of an equilateral triangle and the magnitude of the amino acid sequence similarity between any two species is denoted by a vector that points to the side linking those two species (**Figure 1E**). If each species is equally similar to the other two, the sum of those vectors would place the point at the centroid of the triangle. As can be seen, the data points for TAZ, SAV1, LATS1 and LATS2 are close to the centroid. However, the data point for YAP1 is much closer to the side linking the human to the capybara. This indicates that the human and capybara YAP1 are each more similar to the other than either is to the mouse (YAP1 human to capybara = 99.1%, Human to mouse = 92.8%, and mouse to capybara = 92.5%). If a phylogeny was drawn based solely on YAP1, the capybara and human would be considered sister species with the mouse being the outgroup (**Figure 1F**). This is in contrast to the well-established phylogeny that would place the two rodent species as sister species with the human as the outlier (**Figure 1G**).

### Evidence of relaxed constraint in the YAP1 gene in specific rodent taxa

The neutral theory of molecular evolution proposes that most sequence differences between species are the result of random fixation of mutations (Kimura, 1968, 1986). It is likely, however, that a small subset of these changes are adaptive and have been selected for due to their phenotypic effects. A well-established method for identifying signatures of positive selection in evolution has been to compare the rate of nonsynonymous nucleotide substitutions (*dN*) which, change the encoded amino acid, with the rate of synonymous substitutions (*dS*), which do not (Kimura, 1977; Miyata and Yasunaga, 1980) and test for episodic positive selection across branches of a phylogenetic tree. We performed such an analysis using a codon-substitution model (Goldman and Yang, 1994; Yang, 1998; Yang and Nielsen, 1998; Álvarez-Carretero et al., 2023) in the CODEML package implemented in PAML. This method allowed us to estimate the *dN/dS* ratio (*ω*) for either the entire phylogenetic tree or for portions of it (**Figure 2A-C**). An *ω* of <1 is indicative of purifying selection in a gene where most nonsynonymous mutations are deleterious and eliminated from the population. *ω* = 1 indicates neutral evolution. An *ω* of >1 is indicative of positive selection which is observed in a small subset of genes such as those involved in host-pathogen interactions. For most genes, if the *ω* ratio is averaged across the entire tree and over all sites of the gene, it is typically <1. Indeed, for the phylogeny shown in **Figure 2F**, *ω* = 0.07 indicates that the YAP1 gene is generally under strong purifying selection over deep evolutionary time.

**Figure 2.**
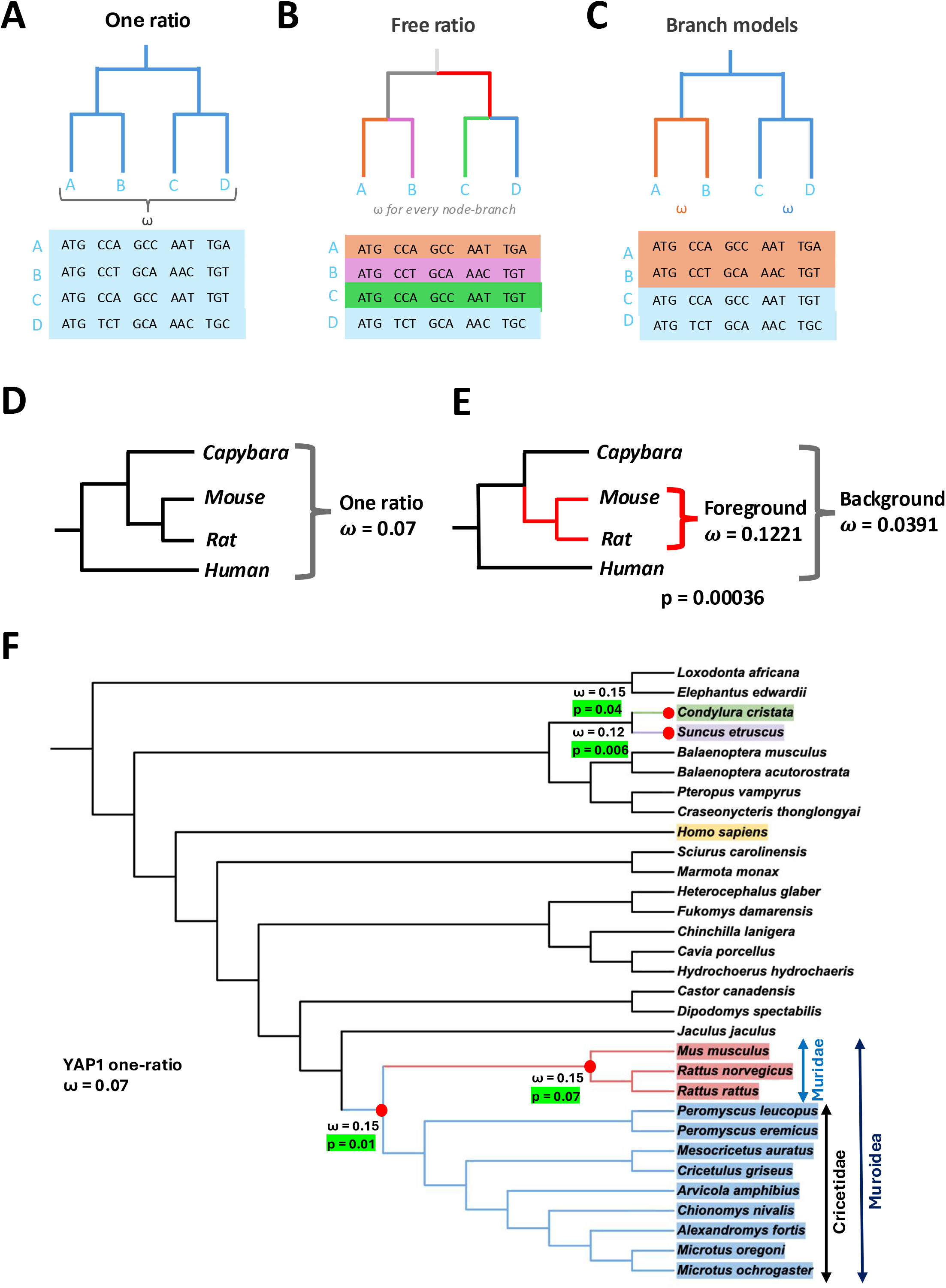
Estimation of dN/dS ratios in the rodent phylogenetic tree. **A-C)** The dN/dS ratio can be averaged across all codons across all species in the tree using a one ratio model **(A)**. Alternate approaches included the free ratio model where an w value is calculated for each node in the phylogeny **(B)**. Branch models **(C)** calculate an *ω* ratio for the foreground branch and compare it to that of the background which is the remainder of the tree (based on Álvarez-Carretero et al., 2023).. **D-E)** Simplified phylogenetic trees composed of three rodent species with the human as outgroup. *ω* is calculated using the one ratio model in **(D)** and the foreground/backgound model in **(E)**. The p-value indicated the likelihood of the null hypothesis, that the *ω* values of the foreground and background do not differ significantly. **F)** The phylogenetic tree used for dN/dS analysis. After computing *ω* values using the foreground/background model, those foreground branches where the *ω* value of the foreground differs from that of the background with a p-value close to or less than 0.05 are indicated. Nodes that were likely under relaxed constraint led to branches consisting of groups Muroidea and Muridae, as well as in the individual species *Suncus etruscus* and *Condylura cristata*.

Since we observed divergence of YAP1 sequences in rodent lineages, we tested the hypothesis that positive selection resulted in a higher rate of nonsynonymous to synonymous substitutions in specific rodent lineages. We first applied this analysis to a simple phylogenetic tree consisting of the human, capybara, mouse and rat. Using the one ratio model which averages *ω* across the entire tree, *ω* = 0.07 (**Figure 2D**). However, when the ancestral node of the mouse and the rat is designated the foreground, the *ω* values for the foreground and background are 0.1221 and 0.0391 respectively with a P=0.00036 (**Figure 2E**). This indicates that in a simple model an ancestor of the mouse and rat exhibits a strong likelihood of relaxed constraint which can be suggestive of positive selection.

To understand more specifically which rodent lineages share YAP1 sequence divergence, we then analyzed 31 species in the placental mammal phylogeny via a two-step approach (Ree et al., 2004). To test the hypothesis that specific branches might be under relaxed constraint and thus undergoing relative positive selection, we first used a free-ratio model to calculate the *ω* value at each node of the phylogenetic tree (**Supplementary Figure S1**). Then, for branches exceeding a cutoff of *ω* > 0.1 we implemented a branch model (Yang, 1998; Yang and Nielsen, 1998) to compare the chosen foreground branch to all the other background branches. From this analysis, four branches show elevated *ω* values relative to background and p-values equal to or less than 0.07 (**Figure 2F**). One branch was at the ancestral node of the Muroidea group, which includes black and brown rats, mice, hamsters and voles (*ω =* 0.15, p = 0.01). Nested within this group were the Muridae, which includes the mouse, the brown rat, and the black rat (*ω* = 0.15, p = 0.07). In addition to these taxa, the two species from the order Eulipotyphla, the star-nosed mole (*ω* = 0.15, p = 0.04) and the Etruscan shrew (*ω* = 0.12, p = 0.006) showed evidence of relaxed constraint.

### Mutations that are predicted to reduce function are found in mouse and vole YAP1

If the changes in the YAP1 protein that have occurred in small rodents have contributed to their size reduction, these changes would be predicted to reduce the activity of YAP1, as it is a protein that promotes the transcription of pro-growth genes. One way to assess the likelihood that these changes deleteriously impact protein function is to use AlphaMissense, a deep learning model that classifies variants in the human proteome (Cheng et al., 2023). The model uses two types of information to calculate the likelihood that a variant is pathogenic, or likely to be deleterious. First, it uses information derived from protein structure predictions, primarily based on AlphaFold. Second, it examines the prevalence of that variant in human and primate populations. Using these criteria, each variant is assigned a pathogenicity score. The cutoffs are based on comparisons to ClinVar labels; scores of < 0.34 are deemed to be likely benign while those > 0.564 are considered likely pathogenic. Scores between 0.34 and 0.56 are considered ambiguous.

We treated the changes found in the capybara, the mouse, and the vole as potentially deleterious mutations in the human protein and classified their likely pathogenicity using AlphaMissense. The capybara, mouse and vole had 11, 42 and 30 differences to the human protein, respectively (**Figure 3A, B**). A small number of variants did not have an AlphaMissense score. However, of those that did, almost all were scored as benign. One difference in the mouse protein, in the transactivation domain, D481E (D481 refers to the corresponding amino acid in humans) was scored as ambiguous. One difference in the vole protein, which maps to the second WW domain, E245D (E245 refers to the corresponding amino acid in humans), is scored as likely pathogenic. Thus, for each of these two small rodent species, there is at least one amino acid change that has the potential to alter protein function. Most other changes are individually considered benign.

**Figure 3.**
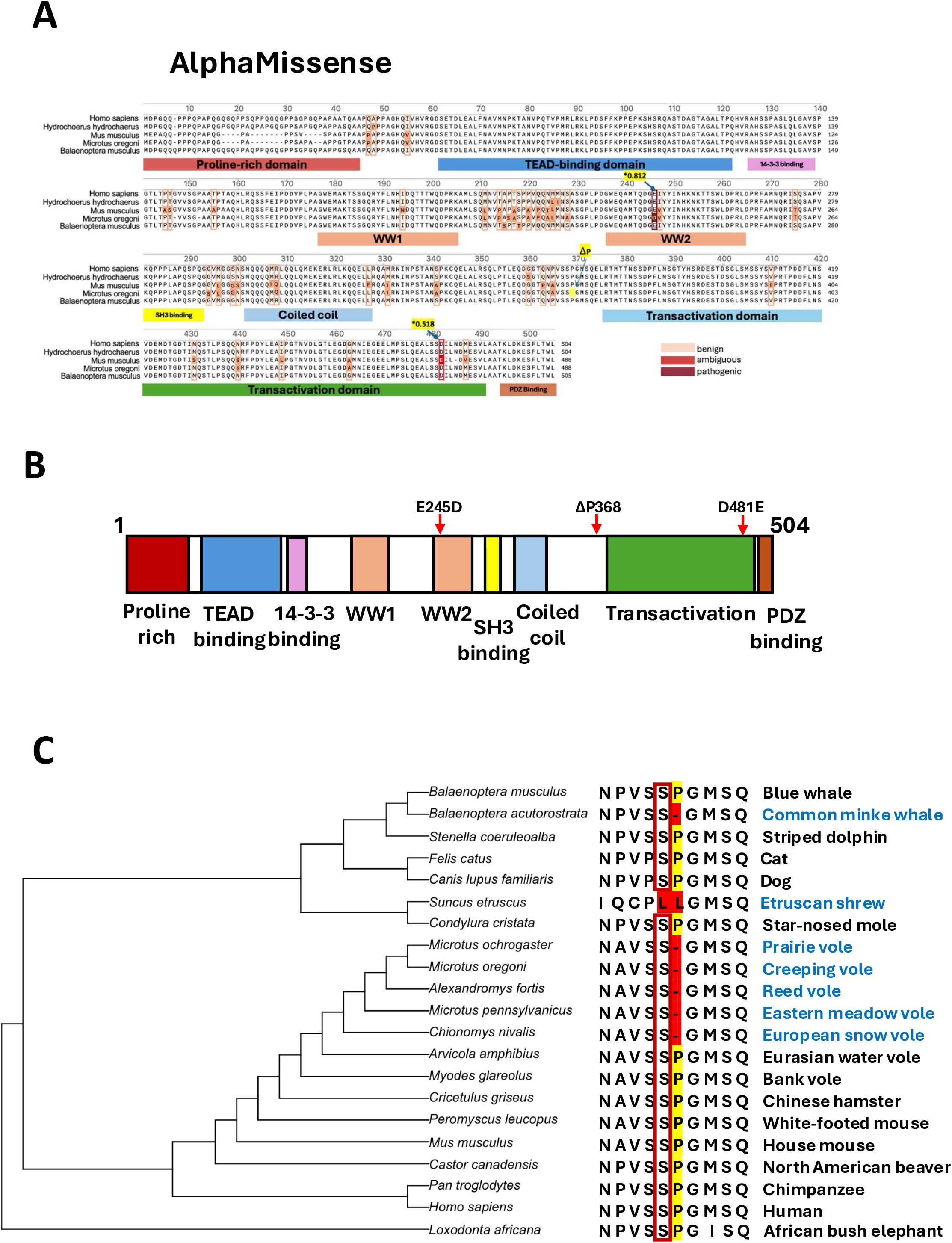
Assessing the impact of coding region changes in YAP1. **A)** Alignment of YAP1 sequences of human, capybara, mouse and vole with the blue whale as the outgroup. Changes in the non-human sequences are treated as potentially pathogenic and their pathogenicity score using AlphaMissense is indicated. **B)** Domains of YAP1, drawn to scale, indicating the positions of the E245D change in the vole protein that was classified as "pathogenic" and the D481E change in the mouse protein that was classified as “ambiguous”. Also shown is the deletion of the highly conserved proline residue, corresponding to P368 in human YAP1, that is deleted in *Microtus oregoni*. **C)** S367 is a site phosphorylated by cyclin-dependent kinases and following ErbB signaling. In this alignment the residues corresponding to S367 are boxed in red. The motif includes P368 which is highlighted in yellow. P368 is deleted in a clade of voles (names in blue) and independently in the Common minke whale, a small whale. Deletions of the proline residue or its substitution are shaded in red. The entire sequence which encodes the phosphorylation motif is absent in the Etruscan shrew.

For the mouse D481E change and the vole E245D change, we used the Suzuki-Gojobori test (Suzuki and Gojobori, 1999) to ask if either of these changes were likely to have occurred as a result of positive selection. This test compares the number of synonymous and non-synonymous substitutions that have occurred from a hypothetical ancestral sequence. Since three of the voles in the phylogenetic tree have the E245D change, they were used as the foreground for the test which showed evidence of positive selection, p = 0.007. The D481E substitution is found in both the mouse and the rat which were used as the foreground. This change did not show evidence of positive selection, p=0.700. These types of analyses do not exclude the possibility that the positive selection occurs on functions of YAP1 that have been altered as a result of multiple amino acid changes.

We also observed that the *Microtus oregoni* YAP1 protein has a deletion of a highly conserved proline residue corresponding to P368 in humans (**Figure 3B**). This deletion is observed in a clade of vole species (**Figure 3C**) but not in all voles. Remarkably, this same proline residue is deleted in the Common minke whale, *Balaenoptera acutorostrata*, which is a relatively small whale species, raising the possibility that this deletion reduces YAP1 activity. The preceding residue, S367 (S352 in the mouse), is phosphorylated (Yang et al., 2013; Aharonov et al., 2020; Uliana et al., 2023). One study implicated CDK1 as the kinase that phosphorylates this site during mitosis which is consistent with the position of the proline following the phosphorylation site (Yang et al., 2013). Another study implicated a kinase downstream of ERBB2, likely ERK, which also requires a proline following the serine, during mouse cardiac regeneration (Aharonov et al., 2020). Importantly, both studies found phosphorylation to enhance YAP1 function. Thus, deletion of the proline would be predicted to reduce YAP1 function. The smallest mammal, the Etruscan shrew, lacks this phosphorylation site altogether (**Figure 3C**). These findings provide support to the notion that small animals, including some outside the rodent clade, have evolved amino acid substitutions in YAP1 that reduce its function.

### YAP1 from different rodent species differ in their growth-promoting activity

Since we have found evidence suggestive of biological consequence in at least two different small rodent species, an attractive hypothesis is that the YAP1 orthologs from small rodents such as the mouse *M. Musculus* and the vole *M. oregoni* have less growth-promoting activity than those from the large rodent, the capybara *H. hydrochaeris* or the human. The quality of sequence in the Etruscan shrew genome was based on fewer reads and thus likely to be less reliable - we therefore did not pursue functional studies on that species and instead focused on mutations seen in the Muridae and Muroidea groups. To assess the growth-promoting activity of human and rodent YAP1 genes in a living organism, we synthesized cDNAs encoding YAP1 for each of these species,including only the coding sequences, and expressed them in the developing *Drosophila* wing (Tripathi and Irvine, 2022) using the GAL4/UAS system (Fischer et al., 1988; Brand and Perrimon, 1993). Expression of either the *Drosophila* YAP1 ortholog *yorkie* (*yki*) or the human ortholog in developing tissues from wild-type *Drosophila* has previously been shown to promote overgrowth (Zhao et al., 2007). This is because this increases the overall levels of *yorkie*/YAP1 in that tissue which result in increased expression of growth-promoting genes.

We therefore generated transgenic *Drosophila* lines that had integrated *UAS-YAP1* (human), *UAS-YAP1* (capybara), *UAS-YAP1* (mouse) and *UAS-YAP1* (vole) at the same genomic locus. Each of these lines was crossed to the *nubbin>GAL4* driver line (*nub>GAL4*) which is expressed in the pouch of the larval wing imaginal disc, the structure that generates the blade of the adult wing (**Figure 4A, B**). Additionally, these larvae carried a copy of a *UAS-GFP* transgene such that the region of YAP1 expression can be easily visualized in the wing disc.

**Figure 4.**
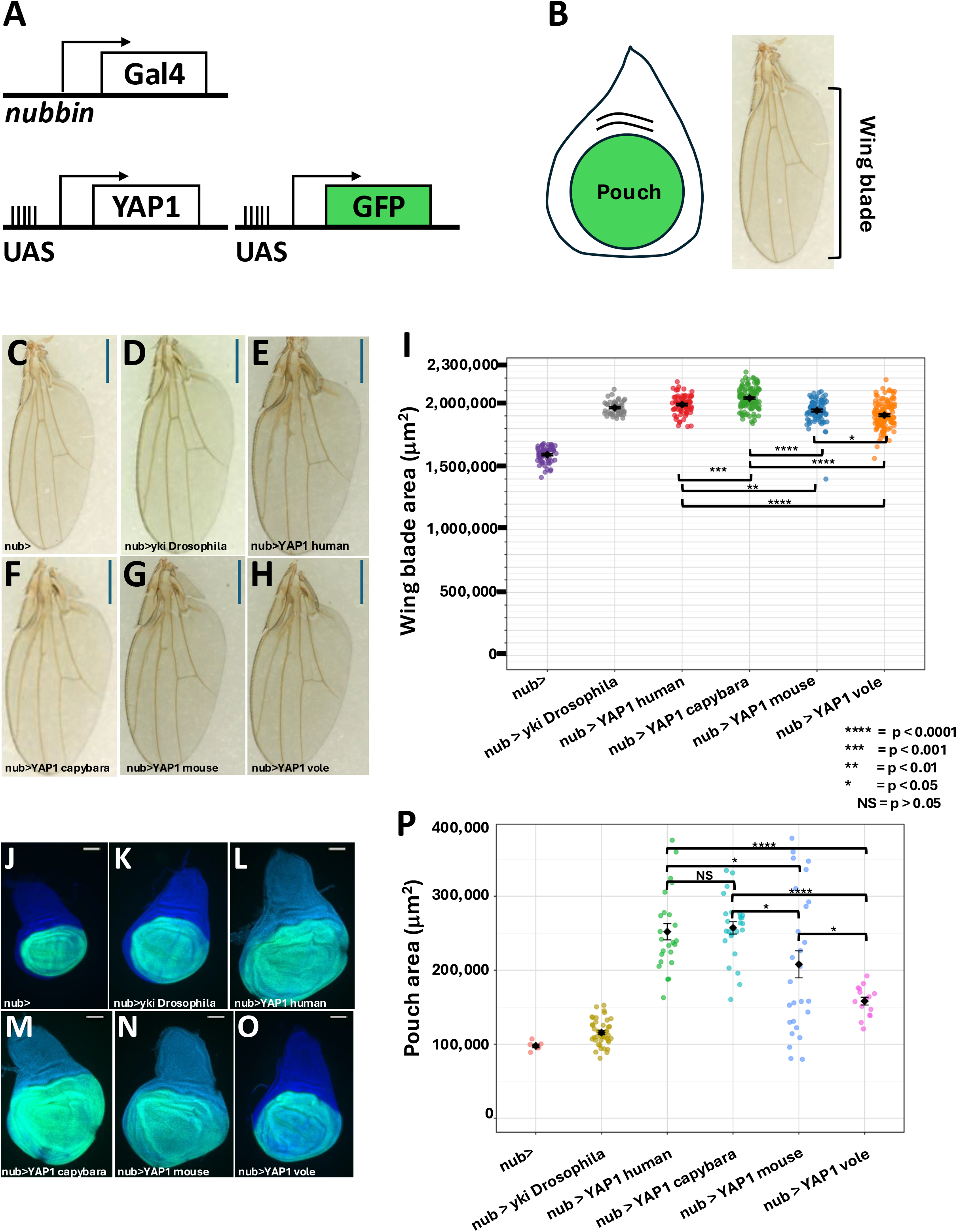
Effect of different YAP1 genes on growth in the wing disc and on adult wing size. **A,B)** The GAL4/UAS system. GAL4 expressed in the pouch of the wing disc under the control of regulatory elements from the *nubbin* (*nub*) gene binds to UAS elements upstream of promoters that express YAP1 as well as GFP. **C-H)** Wings from adult females. **(C)** shows the control that only expresses UAS-GFP. In the remaining samples the following transgenes are expressed **(D)** *Drosophila melanogaster yorkie*, **(E)** human YAP1, **(F)** *Hydrochoerus hydrochaeris* (capybara) YAP1. **(G)** *Mus musculus* (mouse) YAP1 and **(H)** *Microtus oregoni* (vole) YAP1. Wild-type versions of all YAP1 genes were used. Scale bars are 500 μm. **I)** Quantification of data on adult wing blade areas - each sample is shown as a data point. Error bars show standard errors. *nub>* (control), mean=1592648, SD=62446, SE=7931, n=62; *nub>yki* Drosophila, mean=1964685 SD=52120, SE=7446, n=49; *nub>YAP1 human*, mean=1989859, SD=73772, SE=8462; *nub>YAP1 capybara*, mean=2041156, SD=85906, SE=7976.212, n=116; *nub>YAP1 mouse*, mean=1941905, SD=93161, SE=9988, n=87; *nub>YAP1 vole*, mean=1905796, SD=105238, SE=9413, n=125. For all pairwise comparisons using ANOVA and Tukey HSD, p-adj are: *nub>* vs *nub>yki* p=1.795734E-10; *nub>* vs *nub>YAP1(hum)* p=1.795734E-10; *nub>* vs *nub>YAP1 (cap)* p=1.795734E-10; *nub>* vs *nub>YAP1 (mouse)* p=1.795734E-10; *nub>* vs *nub>YAP1 (vole)* p=1.795734E-10; *nub>yki* vs *nub>YAP1 (hum)* p=5.976471E-01; *nub>yki* vs *nub>YAP1 (cap)* p=3.632899E-06; *nub>yki* vs *nub>YAP1 (mouse)* p=6.728666E-01 (not significant); *nub>yki* vs *nub>YAP1 (vole)* p=7.597415E-04; *nub>YAP1 (hum)* vs *nub>YAP1 (cap)* p=8.271747E-04; *nub>YAP1 (hum)* vs *nub>YAP1 (mouse)* p=5.393401E-03; *nub>YAP1 (hum)* vs *nub>YAP1 (vole)* p=8.338463E-10; *nub>YAP1 (cap)* vs *nub>YAP1 (mouse)* p=1.796800E-10; *nub>YAP1 (cap)* vs *nub>YAP1 (vole)* p=1.795735E-10; *nub>YAP1 (mouse)* vs *nub>YAP1 (vole)* p=3.206610E-02. **J-O)** Imaginal discs dissected from female third instar larvae expressing the transgenes. DAPI (blue) stains all nuclei in the disc. GFP expression is confined to the wing pouch. **(J)** shows the control that only expresses *UAS-GFP*. In the remaining samples the following transgenes are expressed **(K)** *Drosophila melanogaster yorkie*, **(L)** human YAP1, **(M)** *Hydrochoerus hydrochaeris* (capybara) YAP1. **(N)** *Mus musculus* (mouse) YAP1 and **(O)** *Microtus oregoni* (vole) YAP1. Wild-type versions of all YAP1 genes were used. Scale bars are 100 μm. **P)** Quantification of data on disc pouch size - each sample is shown as a data point. Error bars show standard errors. Data for each genotype: *nub>* (control), mean=97752, SD=6130.162, SE=2502.628, n=6; *nub>yki Drosophila*, mean=115764, SD=17102.858, SE=2608.161, n=43; *nub>YAP1 human*, mean=251991, SD=53713.259, SE=10964.173, n=25; *nub>YAP1 capybara*, mean=257238, SD=42107.715, SE=8421.543,n=30; *nub>YAP1 mouse*, mean=207962, SD=94900.797, SE=18263.667, n=27; *nub>YAP1 vole*, mean=158203, SD=20312.516, SE=5244.669, n=15. For all pairwise comparisons using ANOVA and Tukey HSD, p-adj are: *nub>* vs *nub>yki* p=9.677781E-01 (not significant); *nub>* vs *nub>YAP1(hum)* p=2.094793E-08; *nub>* vs *nub>YAP1 (cap)* p=5.905382E-09; *nub>* vs *nub>YAP1 (mouse)* p=9.114331E-05; *nub>* vs *nub>YAP1 (vole)* p=1.602056E-01 (not significant); *nub>yki* vs *nub>YAP1 (hum)* p=3.930190E-14; *nub>yki* vs *nub>YAP1 (cap)* p=2.631229E-14; *nub>yki* vs *nub>YAP1 (mouse)* p=4.953161E-10; *nub>yki* vs *nub>YAP1 (vole)* p=7.689847E-02 (not significant); *nub>YAP1 (hum)* vs *nub>YAP1 (cap)* p=9.992592E-01 (not significant); *nub>YAP1 (hum)* vs *nub>YAP1 (mouse)* p=3.488065E-02; *nub>YAP1 (hum)* vs *nub>YAP1 (vole)* p=2.911063E-06; *nub>YAP1 (cap)* vs *nub>YAP1 (mouse)* p=1.057946E-02; *nub>YAP1 (cap)* vs *nub>YAP1 (vole)* p=5.587449E-07; *nub>YAP1 (mouse)* vs *nub>YAP1 (vole)* p=3.974131E-02.

We first measured the area of the adult wing blade from each genotype (**Figure 4C-I**). Since wings from males and females differ in size, we measured only female wings, and we measured an average of 109 wings for each of the three rodent species. The area of adult wings from *nub>YAP1 capybara* were 5% larger than those of *nub>YAP1 mouse* (p=1.8E-10) and 7% larger than *nub>YAP1 vole* (p=1.8E-10). The *nub>YAP1 vole* wings were 2% smaller than those of *nub>YAP1 mouse* (p=3.2E-2). The overgrowth elicited by *UAS-YAP1 human* is between the capybara and mouse genes. *UAS-yki Drosophila* and the line expressing the nub-GAL4 driver alone are also shown for comparison (Statistical data for all genotypes are shown in the Figure Legend).

We also dissected wing imaginal discs from staged female third-instar larvae and measured the area of the wing pouch which generates the adult wing blade. This was done by measuring the size of the GFP-expressing domain from each genotype (**Figure 4J-P**). Wing pouch sizes show more variability when compared to adult wing sizes, firstly because of small differences in larval maturity that exist while disc is still growing, and secondly due to the wing pouch of the disc is not as flat as the adult wing blade. Despite these caveats, the pouch sizes from the four genotypes are ranked in the same order as was observed with the adult wings. Pouches expressing *nub>YAP1 mouse* are 19% smaller than those expressing *nub>YAP1 capybara* (p=1.1E-2) and those of *nub>YAP1* vole are 38% smaller than those expressing *nub>YAP1 capybara* (P=5.6E-7). The *nub>YAP1* vole pouches were 2% smaller than those of *nub>YAP1 mouse* (p=4.0E-2). Once again, growth elicited by *nub>YAP1 human* is between that of the *YAP1 capybara* and *YAP1 mouse*. The differences in pouch size exceed those observed in the adult wing possibly because there are compensatory effects during the pupal stage of development. In any case, both adult wing measurements and the sizes of larval wing disc pouches show that the YAP1 genes from species from two different taxa of small rodents, the Muroidea and the Cricetidae, have less growth promoting activity than the large rodent, the capybara.

### Mammalian YAP1 genes activate expression of known target genes of *Drosophila yki*

If the mammalian genes promote growth in *Drosophila* wing discs in a similar way to *Drosophila yki*, it is likely that they are capable of activating known target genes of *Drosophila yki*. One well characterized target is *diap1,* which encodes the antiapoptotic protein Diap1. Elevated expression of Diap1 protein is observed in mutations that are known to increase nuclear *yki* activity (Tapon et al., 2002). To determine whether the different YAP1 transgenes were capable of increasing *diap1* expression, we expressed each transgene as well as GFP in the posterior compartment of the wing disc using the *hh-GAL4* driver line (**Figure 5 A-F**’). When stained with anti-Diap1 antibody, expression of Diap1 protein was elevated in the posterior compartment of discs with each of the mammalian YAP1 genes. Increased Diap1 expression was also observed with *Drosophila yki*, although its expression was not as elevated as with the mammalian genes. This is consistent with the increased growth elicited by the mammalian YAP1 genes when compared to *Drosophila yki*. Thus, as has been shown previously for human YAP1 (Zhao et al., 2007), each of the mammalian YAP1 genes tested is capable of activating *diap1* expression in *Drosophila* imaginal discs. However, the uneven expression of Diap1, at least as detected by the antibody, made it difficult to reliably compare the effect of the different rodent genes. We therefore used an assay in cultured mammalian cells (see below).

**Figure 5.**
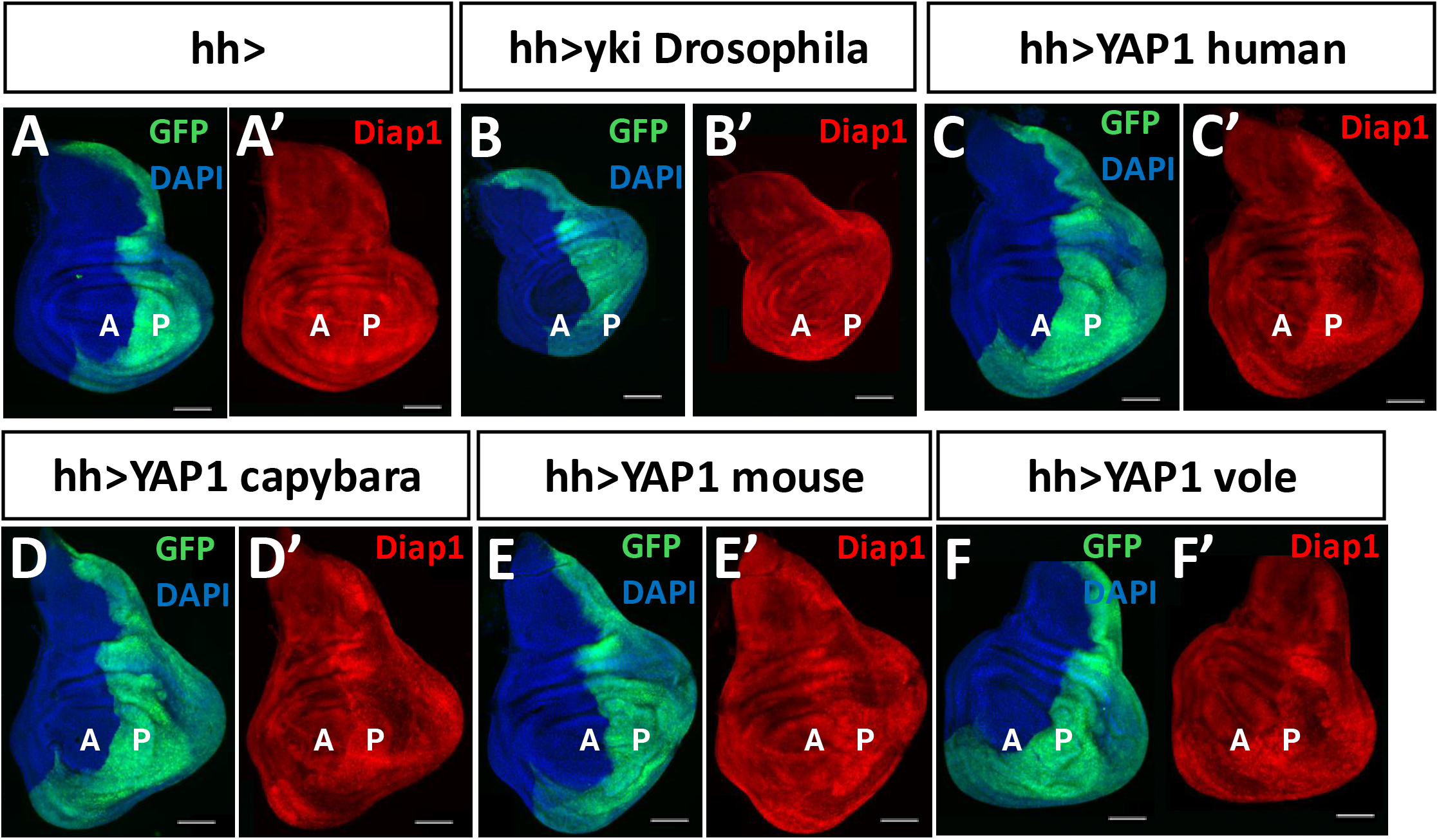
Expression of mammalian YAP1 proteins in *Drosophila* wing discs upregulates Diap1 levels. **A-F)** Wing imaginal discs dissected from third instar larvae. Expression of transgenes expressed under the control of UAS elements driven by *hedgehog-GAL4* (*hh-GAL4*). All larvae also carry a *UAS-GFP* transgene to demonstrate the posterior compartment (P) where *hh-GAL4* is expressed and the anterior compartment (A) where it is not. **(A)** and **(A’)** show control discs that do not carry a *UAS-YAP* transgene. All others carry either *Drosophila* yki **(B, B’)** or one of the mammalian YAP1 genes **(C-F, C’-F’)**. Scale bars are 100 μm.

### YAP1 proteins from the mouse and vole show reduced TEAD-dependent transcriptional activity compared with those from human and capybara in mammalian cells

YAP1 does not bind DNA directly. Instead, it functions as a transcriptional co-activator by interacting with DNA-binding transcription factors, most prominently TEAD family proteins, to activate downstream target genes (Zhao et al., 2008). We therefore compared the transcriptional activity of YAP1 proteins from the four species examined in this study using a TEAD-responsive luciferase reporter assay (**Figure 6A, B**). HEK293T cells were transfected with FLAG-tagged YAP1 genes from human, capybara, mouse and vole together with a luciferase reporter driven by a synthetic promoter containing multiple TEAD-binding sites. To account for the differences in YAP1 protein abundance among the four constructs, luciferase activity was normalized to YAP1 protein expression. Lysates from three independent luciferase experiments were pooled separately for each of the four YAP1 constructs and analyzed by Western blotting with anti-FLAG and anti-GAPDH antibodies. FLAG-YAP1 band intensities were quantified relative to GAPDH. The human YAP1/GAPDH ratio was set to 1.0 and the YAP1/GAPDH ratios for the other constructs were expressed relative to the human value. As shown in **Figure 6B**, the resulting relative ratios were 1.00 for human, 1.50 for mouse, 1.25 for capybara, and 2.00 for vole YAP1. The Luciferase activity for each YAP1 construct was then normalized to the corresponding relative FLAG-YAP1/GAPDH value (**Figure 6A**).

**Figure 6.**
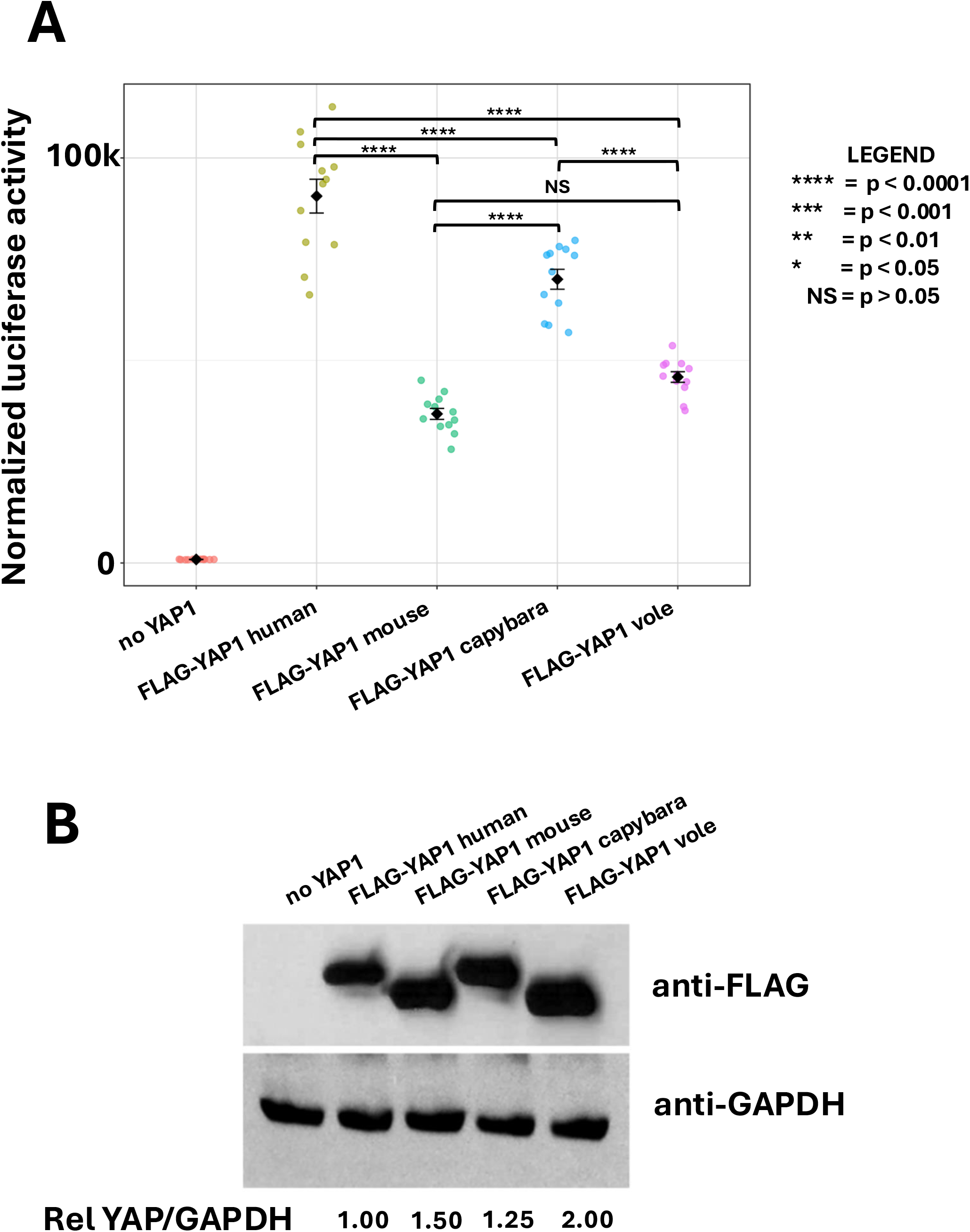
Activation of a TEAD-dependent luciferase reporter in mammalian cells by YAP1 proteins from different species. **A)** Luciferase activity for each YAP1 construct after normalization to the corresponding relative FLAG-YAP1/GAPDH level. Values were divided by the relative FLAG-YAP1/GAPDH ratio shown in **(B)**. Normalization was not applied to the “ no YAP1” control. Each point represents one replicate transfection. Four replicate transfections were performed in each of three independent experiments (12 measurements per construct). Error bars indicate SE. Values: No YAP1 mean=872.08, SD=54.3, SE=15.67, n=12; human mean= 90571.67, SD=14457.8, SE=4173.61, n=12; mouse mean=36802.8, SD=4698.2, SE=2034.38, n=12; capybara mean=70046.0 SD=8519.7, SE=3074.29, n=12; vole mean=45906.2, SD=4539.5, SE=2620.9, n=12. For pairwise comparisons using ANOVA and Tukey HSD p-adj: no YAP1 vs human p=3.610334E-12; no YAP1 vs mouse p=3.659184E-12; no YAP1 vs capybara p=3.610334E-12; no YAP1 vs vole p=3.611667E-12; human vs mouse p=3.610334E-12; human vs capybara p=6.385669E-07; human vs vole p=3.611889E-12; mouse vs capybara p=4.026446E-12; mouse vs vole p=5.670476E-02 (not significant); capybara vs vole p=1.028383E-08. **B)** Western blot analysis of pooled lysates from HEK293T cells transfected with the TEAD-responsive luciferase reporter with or without different FLAG-tagged YAP1 constructs from human, mouse, capybara, or vole. FLAG-YAP1 band intensities were quantified by densitometry and normalized to the corresponding GAPDH signal The human FLAG-YAP1/GAPDH ratio was set to 1.0 and the ratios for the other YAP1 constructs were expressed relative to the human value. Densitometric values were: human YAP1=261031, GAPDH=213717, ratio=1.22, relative ratio set to 1.0; mouse YAP1=396376, GAPDH=216599, ratio=1.83, relative ratio = 1.5; capybara YAP1=313345, GAPDH=204800, ratio=1.53, relative ratio = 1.25; vole YAP1=517346, GAPDH=212027, ratio=2.44, relative ratio = 2.

As shown in **Figure 6A**, human YAP1 exhibited the highest reporter activity. Capybara YAP1 had higher activity than mouse YAP1 (p=4.0E-12) and vole YAP1 (p=1.0E-08). The activity of the mouse and vole YAP1 was similar (p=5.7E-02). Thus consistent with their reduced growth-promoting properties in the developing *Drosophila* wing, mouse and vole YAP1 show reduced TEAD-dependent transcriptional co--activator activity compared to human and capybara YAP1.

## DISCUSSION

Our study provides evidence that coding-region changes in the YAP1 gene could contribute to size differences between rodent species. We provide five lines of evidence. First we show that while YAP1 is highly conserved between diverse mammals such as humans, blue whales and elephants, it has diverged considerably among smaller rodent species. Strikingly, the human and capybara proteins are more similar to each other than either is to the mouse protein. Second, by calculating the ratio of synonymous to non-synonymous substitutions, we find evidence for relaxed constraint in what is otherwise a highly conserved protein in two clades of rodents - the Muroidea, and within it, the Muridae. Third, the YAP1 proteins of the mouse and vole have amino acid changes that would be consistent with alterations in their activity. Fourth, when expressed in the developing *Drosophila* wing, the mouse and vole YAP1 have less growth-promoting activity than human or capybara YAP1. Finally, when transfected into mammalian cells, the mouse and vole YAP1 are less effective at promoting transcription, suggesting that they would be less effective at activating the expression of growth-promoting target genes under physiological conditions.

Our findings do not imply that the size differences between the capybara and the small rodent species are entirely, or even mostly, due to differences in YAP1. There are at least six different pathways that have been shown to regulate growth even in *Drosophila* imaginal discs (Hariharan, 2015) and there are likely to be many more such pathways in complex vertebrates. Moreover, changes in one or more other pathways acquired at similar times in the evolutionary history of the species, as those that occurred in YAP1, could either enhance or suppress the effect of changes in YAP1. Additionally, changes in different components of the Hippo pathway could also occur at different points in evolution. The increasing availability of large numbers of sequenced genomes should facilitate comparisons of the coding region differences between large numbers of genes that function in growth regulation between any two species. Such studies will also illuminate how frequent coding region changes in growth-regulatory genes between members of the same species are and how representative the reference genomes that we have used are for that species as a whole. What is likely to be more difficult and labor intensive is the assessment of the functional consequences of those changes, as we have attempted to do in this study.

Our results do not preclude the possibility that changes in expression levels or in the expression patterns of these same genes also have a role in generating size differences between different species. Indeed it has been argued that most morphological evolution is driven by changes in cis-regulatory regions because changes in coding regions of genes that are highly conserved are more likely to be deleterious, especially if those gene products function to regulate many different aspects of biology (Carroll, 2008). Work in the last decade has shown that YAP1 does indeed function in a large variety of biological processes in different species including growth regulation, cell fate determination, and the regulation of cytoskeletal dynamics (Zheng and Pan, 2026). Consistent with the notion that most changes in such a gene might be expected to be deleterious, YAP1 is highly conserved between different species. However, experimental studies have shown that manipulation of YAP1 levels can cause changes in organ size (Camargo et al., 2007; Dong et al., 2007). Presumably a similar effect can be achieved by mutations whose effect is to preserve all functionalities of a protein, yet change the level of its activity, possibly in subtle ways. Perhaps this is why there are many changes between the small and large rodents where each change, in itself, has a very subtle effect on protein activity.

Finally, we have demonstrated the utility of the *Drosophila* wing imaginal disc as an in vivo assay for the growth-promoting activity of YAP1 genes from different species. The obvious advantage of this assay is the ease of experimental manipulation and the rapid generation time of *Drosophila*. We have attempted to complement our results with the measurement of the transcriptional activation properties of the different YAP1 proteins in mammalian cells in culture. These results provide the impetus for more long-term studies where the YAP1 genes between different rodent species could be swapped, which would ensure that those genes are expressed at near-physiological levels and in appropriate spatiotemporal patterns in a related species.

## METHODS

### Sequence alignments and comparisons

YAP1, TAZ, SAV1, LATS1, and LATS2 sequences were obtained from NCBI. Sequence .fasta files and method of isoform trimming is described in the Hariharan Lab Github under Bansal_Hariharan/fig.1/sequence_files. Because many gene sequences were available per gene, UniProt was used to ensure the same isoforms were used in each case (Ahmad et al., 2025). For **Figure 1C**, COBALT was used to align *Homo sapiens*, *Hydrochoerus hydrochaerus*, *Mus musculus*, *Microtus oregoni*, and *Balaenoptera musculus* YAP1 (Papadopoulos and Agarwala, 2007).

### Sequence matrix

Protein pairwise matrices for **Figure 1D** were produced by first creating a trimmed alignment for ∼30 rodent species; sequences were trimmed using Gblocks (Castresana, 2000; Talavera and Castresana, 2007) and aligned using Clustal Omega (Sievers et al., 2011). Gblocks trimming used the following default parameters: Minimum number of sequences for a conserved position: 50% of the total sequences + 1. Gap cutoff: None allowed. Maximum number of contiguous nonconserved positions: 8. Minimum length of a block: 10. Minimum number of sequences for a flank position: 85% of the total sequences.

NGPhylogeny.fr was primarily used as a web-based workflow for sequence cleaning/trimming and alignment, as it provides a comprehensively usable workflow (Lemoine et al., 2019). Cleaned alignments were then fed into a Python script used to parse the alignment one amino acid at a time and assign amino acid identity scores, then outputting a matrix with each species’ pairwise score. Identity was defined only by an exact match of amino acids. The Python script, raw sequences, cleaned sequences, and output matrices are also on Github under Bansal_Hariharan/fig.1/All_pairwise_matrix_files.

### Species’ average mass

The average mass of placental mammalian species for **Figure 1D** was derived from the following databases: AnAge was primarily used to obtain average mass (de Magalhães et al., 2024). For species with information that was either missing or in need of corroboration from AnAge, we used the ADW for *Craseonycteris thongolai* (Burns, 2023), Treatment Bank for *Alexandromys fortis*, and *Chionomys nivalis* (Wilson et al., 2017a, 2017b), and MammalBase for *Microtus oregoni* (Lintulaakso, 2021; Wilson et al., 2005).

### Phylogenetic trees

To infer general phylogenetic relationships, a standard placental mammalian phylogeny was used (Christmas et al., 2023). Certain phylogenetic relationships, particularly for the vole species, are still contested. Multiple sources were consulted to ensure usage of the most accurate possible vole phylogeny for the purposes of dN/dS analyses. (Bradley et al., 2004; Robovský et al., 2008; Blanga-Kanfi et al., 2009; Fabre et al., 2012; Herrera-Álvarez et al., 2021). To produce visual trees, simple R code using the library “ape” was used as well as the web-based iTOL (Letunic and Bork, 2024).

### Barycentric plot

Barycentric plots in **Figure 1E** were produced using an R script that created vector weights and assigned weighted averages to each gene based on sequence similarity between the three species. The weighted vector was placed on a particular position in the graph based on sequence identity. The code for this is also available on the lab GitHub under Bansal_Hariharan/fig.1/Barycentric.

#### Calculation of dN/dS ratios

##### Sequence preparation

Codon-aware alignments and trimming of poorly aligned regions were performed using PRANK and BMGE 1.12_1, respectively (Criscuolo and Gribaldo, 2010; Löytynoja, 2014). Both of these tools were used within the NGPhylogeny.fr workflow (Lemoine et al., 2019).

When using BMGE to trim the sequences, specific parameters were used to prevent over-trimming that would remove too much valuable sequence data. These parameters were: Entropy cutoff: 0.5. Gap cutoff: 0.8. Sliding Window Size: 3. Block size: 1.

Alignments were produced within NGPhylogeny.fr in a .phylip format as well as in .nwk format for further downstream analyses (Guindon et al., 2010; Junier and Zdobnov, 2010; Lemoine et al., 2018).

##### Analysis

PAML’s CODEML was used in the command-line to analyze the placental mammalian sequences for evidence of positive selection (Goldman and Yang, 1994; Yang, 1998; Yang and Nielsen, 1998; Álvarez-Carretero et al., 2023). An LRT (likelihood ratio test) and chi-square test were used to assess significance of the branch model results against the one-ratio. The simple code for performing these tests, the .phylip file, the .nwk files, and the PAML CODEML outputs are all in the GitHub under Bansal_Hariharan/fig.2/YAP1.

Note that we performed both free-ratio and branch tests. The branch test, which takes a foreground and background branch as input for the software, will indicate if the omega value for the foreground branch is higher than that of the background branch. This is considered a more robust test than the free-ratio analysis due to the usage of fewer parameters (Ree et al., 2004). Hence, the nodes of interest indicated by the free-ratio test were used to indicate which branches to single out for branch testing.

The CODEML rst1 output also included Suzuki-Gojobori results, which were used in the YAP1 analysis to indicate whether the mutations in *Mus musculus* and *Microtus oregoni* flagged as pathogenic or ambiguous by AlphaMissense were likely to have occurred as a result of positive selection (Suzuki and Gojobori, 1999). Each rst1 file is under Github Bansal_Hariharan/fig.2/YAP1.

#### Alpha Missense

AlphaMissense (Cheng et al., 2023) was used to retrieve pathogenicity scores for mutations in the YAP1 gene seen in other organisms. The web-based AlphaMissense was used.

#### DNA constructs and generation of transgenics

For transfection into mammalian cells, cDNAs for the human, *Hydrochoerus hydrochaerus*, *Mus musculus*, and *Microtus oregoni* were synthesized by Genscript and cloned into the pCDNA3.1 N-DYK vector that expresses each YAP protein (codon 2 onwards) in frame to an N-terminal FLAG epitope tag. For generation of transgenic *Drosophila* lines, the YAP cDNA without the FLAG tag was amplified from the pCDNA3.1 N-DYK construct using a 5’ primer that added a Kozak consensus sequence and an ATG start codon and cloned into the pUASt-attB vector. All constructs were verified by sequencing. Transgenic lines were generated by Bestgene using PhiC31 recombinase and integrated into the attP1 site at 55C4 or chromosome 2. Homozygous stocks were generated for all transgenic lines and subsequently used for the crosses described. The *UAS-yki* stock used was *UAS-ykiV5.0* (Bloomington Stock # 28819).

#### Immunohistochemistry and microscopy

Wing discs were dissected from wandering third instar larvae. To reduce the confounding effect of size differences between sexes, all wing discs were dissected from female larvae and processed as described by Spitzer et al., 2025. The mouse anti-Diap1, a gift of Bruce Hay (Caltech) was used at a concentration of 1:200. The secondary antibody used was goat anti-mouse IgG Alexa Fluor 555 (concentration of 1:500 Thermo Fisher Scientific A-21422, RRID:AB_2535844). Nuclei were stained with DAPI (1:1000, Cell Signaling). Discs were mounted on microscope slides using Invitrogen mountant and imaged with a Zeiss Axio Imager M2 equipped with a 10x objective (for imaging of nub>GFP discs), LED light, AxioCam 506 mono camera, and ApoTome.2 slider for optical sectioning. Images were digitally acquired with the ZEN 2.3 software and processed using FIJI software (Schindelin et al., 2012). All nub>GFP and DIAP1 images have been MAX projected. Pouch area was measured using the Analyze>Measure function on FIJI and image metadata was consulted to ensure that the same set scale was used for the 100 μm scale bar.

For DIAP1 antibody stained discs a 20x magnification to achieve a high enough resolution to visualize cell staining. Some discs were quite large and multiple apotome photos were taken and later stitched together using the Plugins>Stitching>Pairwise Stitching function on FIJI.

Adult fly wings were dissected from relatively young female flies and mounted on Gary’s Magic Mountant (Lawrence et al., 1986) and imaged with a Keyence VHX-5000 microscope, using the 20-200× lens at 100x. A 500 μm scale bar was added during the imaging process. Images used were taken on different days and display slightly varying levels of brightness and contrast. Adult wing area was measured by displaying the image on FIJI, tracing the area from a fixed point at the junction of the wing blade and the hinge and then using the Analyze>Measure function to retrieve wing area.

For qualitative comparisons of wing sizes, wing images or traced silhouettes were overlaid in Microsoft PowerPoint. For quantitative comparisons of wing sizes, wings were traced and area was quantified in FIJI. Charts were generated using the ggplot2 package in Rstudio (Wickham, 2016).

#### Luciferase assays and Western blotting

HEK293T cells were cultured in Dulbecco’s modified Eagle’s medium (DMEM; cat. #11995-065, Gibco) with 10% fetal bovine serum (FBS, BenchMark#100-106, GeminiBio) plus 1% Penicillin-Streptomycin (100x, cat. #15140-122, Gibco).

For measurement of YAP transcriptional activity, a total of 2.5 μg DNA (including 100 ng of 8xGT-IIC-δ51LucII Luciferase reporter construct and/or 400 ng of the indicated FLAG-YAP plasmids) was transiently transfected into HEK293T cells by Lipofectamine 3000 Reagent (cat. #L3000008, Invitrogen/ThermoFisher Scientific) according to the manufacturer’s instructions. The luciferase activity was measured at 36h after transfection and Western blotting were performed as described previously (Zhu et al., 2007). For the Western blot, the corresponding lysates from three independent luciferase experiments (total of 12 human YAP lysates, 12 mouse YAP lysates, 12 capybara YAP lysates, and 12 vole YAP lysates), were pooled and run on the same Western blot. Flag-YAP signal was normalized to GAPDH, with hYAP/GADPH set to 1.0. Values are indicated in the figure legend.Antibodies used were: anti-Flag (M2; cat. #F1804), from Sigma-Aldrich (St. Louis, MO); anti-GAPDH (cat. #sc-47724), from Santa Cruz Biotechnology (Dallas, TX).

## Acknowledgements

We thank Bruce Hay for the gift of the anti-Diap1 antibody. We thank Rasmus Nielsen and John Heulsenbeck for guidance on performing maximum-likelihood modeling for phylogenetic analysis. We thank William Sun, Joyner Cruz, and Zohra Allata for advice and help with experiments. We thank Priya Moorjani, Rasmus Nielsen, David Weisblat, and Noah Whiteman for comments on the manuscript.

---

**Supplementary Figure S1.**
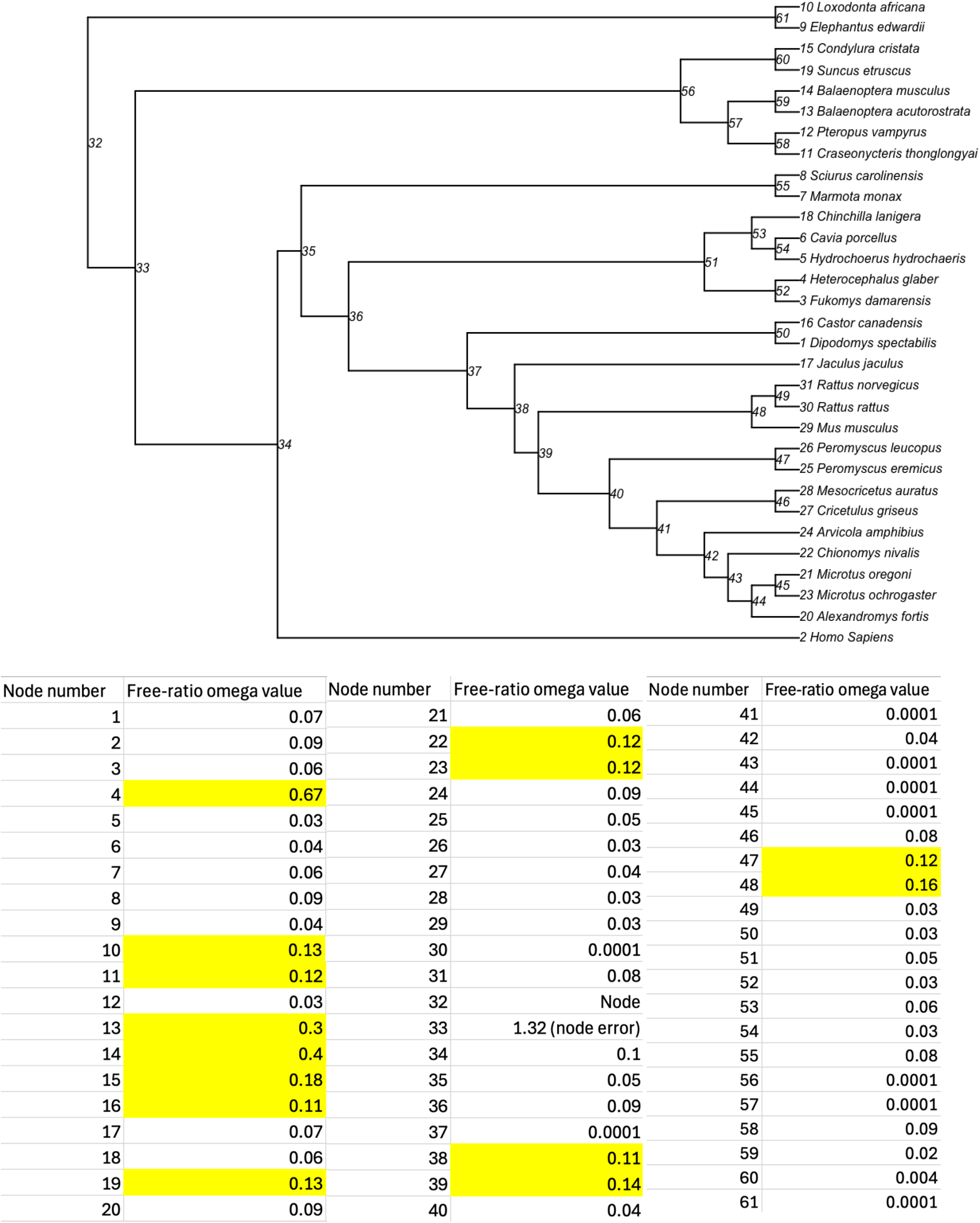
Free ratio ω values for all nodes in the phylogenetic tree. Values that were >0.10 (highlighted) were examined further using the foreground-background model.

